# Behavior choices amongst grooming, feeding, and courting in *Drosophila* show contextual flexibility, not an absolute hierarchy of needs

**DOI:** 10.1101/2025.05.09.653186

**Authors:** Carla E Ladd, Julie H Simpson

## Abstract

To determine the algorithmic rules and neural circuits controlling selection amongst competing behaviors, we established assays where adult *Drosophila melanogaster* choose between grooming and feeding, grooming and courting, or feeding and courting. We find that there is not an absolute hierarchy: while flies typically perform grooming first, they can choose to feed if sufficiently starved, or court if an appropriate female is available. Flies alternate between competing behaviors, performing short bouts of each action rather than completely satisfying one drive before transitioning to another. While we did not do an exhaustive screen, from the candidates we examined, we did not find evidence for a specific genetic or neuronal locus that affects all decisions. We did identify genetic background effects, suggesting that multiple genes may contribute to decision-making priorities. Our results add to a growing body of work on decision-making in *Drosophila* and provide a foundation for future investigation of the exact neural circuits required to achieve appropriate choices.

**Summary Statement:** Flies alternate between competing behaviors. The choice amongst positive actions depends on context and relative drive levels. No central decision-making locus was identified.

## Introduction

In 1943, American psychologist Abraham Maslow published *A Theory of Human Motivation* in which he proposed what is commonly referred to as “Maslow’s Hierarchy of Needs” (Maslow, 1943). Later depicted as a pyramid, this theory states that there are five universal human needs. They rank, from strongest, highest priority to weakest, lowest priority: physiological, safety, love, esteem and self-actualization. The implication is that when needs compete, behaviors that satisfy them will be chosen in priority order. This hierarchy serves as our framework for studying *Drosophila* decision-making in these new choice-based behavior assays.

Decision-making impairment can be a symptom of human neurological conditions including Parkinson’s Disease (Brand et al., 2004; Ryterska et al., 2014), Alcohol Use Disorder (Galandra et al., 2020), Major Depressive Disorder (Rinaldi et al., 2019), Conduct Disorder (Salvatore & Dick, 2018), schizophrenia (Barry et al., 2020), and drug, alcohol, and gambling addictions (Mizoguchi & Yamada, 2014; Lawrence et al., 2009). Human decision-making has been studied using behavioral assays such as Gambling Tests (Bachara et al., 1994; Rogers et al., 1999), the Dictator Game (Knafo et al., 2008), and the Information Sampling Test (Clark et al., 2006). A better understanding of decision-making mechanisms might lead to improved therapies.

Animal models have contributed to decision-making research. In primates, the decision-making process has been studied using mock market economies (Lakshminarayanan et al., 2010). In rodents, decision-making is studied using techniques such as cost-benefit analysis (Alabi et al., 2019; Floresco et al., 2008) and judgment of visual stimuli (Odoemene et al., 2018). Choices between feeding and parenting (Alcantara et al., 2025), feeding and safety (de Araujo Salgado et al., 2023), and prioritization of feeding and courtship (Burnett et al., 2019) have been investigated behaviorally and neuronally in mice. In birds, decision-making has been studied through field observations (Rodríguez et al., 2001). Relevant neural circuits have been investigated in pond snails (Pirger et al., 2014), sea slugs (Gillette & Brown, 2015) and leeches (Briggman et al., 2005; Gaudry & Kristan, 2009). The comparison of neural circuits and activity dynamics that underlie decision-making in different animals and contexts has revealed the complexity of this process; many open questions remain.

We were interested in whether there is a common neural locus that contributes to decision-making more generally, and what algorithms provide flexibility when all of the choices are beneficial. *Drosophila melanogaster* has been an advantageous model in which to study the genes, neurons, and circuits that contribute to perceptual decisions (DasGupta et al., 2014; Groschner et al., 2018), food selection (Münch et al., 2022), mate choice (Hindmarsh Sten et al., 2025; Dickson, 2008), mating duration (Gautham et al., 2024), oviposition site preference (Joseph et al., 2009; Yang et al., 2008), feeding and courtship (Cheriyamkunnel et al., 2021), feeding and locomotion (Mann et al., 2013), and among escape strategies (von Reyn et al., 2014; von Reyn et al., 2017; Asinof and Card, 2024). Often these assays examine the trade-off between benefit and cost: will a male fly continue to mate in the presence of a threat (Cazale-Debat et al., 2024)? Do larvae risk hypoxia for food (Kim et al., 2017), or accept cold or bitter food, if they are very hungry (Wu et al., 2005)? Will they stay on a mediocre but available food source or seek richer resources (LeDue et al., 2016; de Belle et al., 1989)? Switching amongst individual and social behaviors driven by internal mechanisms has also been assayed (Berman et al., 2016).

While Maslow’s theory suggests a more rigid hierarchy of priorities, flexibility in decision-making is a hallmark of successful species (Devineni and Scaplen, 2021). The best choice can change with circumstances, and an animal that can adjust its actions according to risks and rewards, strengths of competing drives, and available resources, thrives in a range of environments. Flies use contextual cues to make behavior choices: flies make decisions about feeding based on how hungry they are, what specific nutrients they lack, what food is available, and prior experience (Nhuchhen et al., 2025; Marella et al., 2012). *Drosophila* can adjust meal size, select sugar or protein-rich foods, and will accept bitter or cold but nutritious substances if sufficiently motivated (LeDue et al., 2016; de Belle et al., 1989). Fruit flies can also avoid food that previously made them sick (Stensmyr et al., 2012). Genes, neurons, and chemical cues that affect feeding decisions include *leucokinin* (Yurgel et al., 2019), *dopamine* (Jin et al., 2023), *diuretic hormone 31* (Lin et al., 2022), and *tyramine* (Cheriyamkunnel et al., 2021).

Flies also make decisions about courtship and mating actions. A male fly will evaluate whether a potential partner belongs to the same species, is male or female, and is mated or virgin (von Philipsborn et al., 2023). Once courtship is initiated, male flies can escalate from orientation and chasing to wing singing depending on the cues from the female fly, and may then proceed to tapping, licking, and attempted copulation (Villella and Hall, 2008). The commitment to copulation and its duration can be influenced by environmental threats: males adjust their choice between courtship and escape depending on external sensory cues, internal drives, and past experiences (Simon et al., 2011). Neural and genetic bases include *leucokinin* (Chen et al., 2025), *tyramine* (Cheriyamkunnel et al., 2021), and *fruitless* (Yamamoto and Koganezawa, 2013).

Our lab’s analysis of grooming behavior indicated that choice of which body part to clean is driven both by the acute distribution of debris and an intrinsic set of priorities: the eyes, antenna, and head are cleaned first (Seeds et al., 2014) – unless there is much more stimulation to mechanosensory bristles in posterior regions of the body (Zhang et al., 2020). An innate hierarchy organizes the grooming sequence from anterior to posterior body parts, with contextual flexibility to respond to the current sensory load. Neural circuits that contribute to the grooming sequence include dopaminergic (Pitmon et al., 2016) and descending neurons (Guo et al., 2022).

Feeding, courting, and grooming are complex motor sequences made up of choices among smaller actions. We noticed, anecdotally, that grooming behavior superseded escape, courtship, and feeding in our standard grooming assays and we wondered if there was an absolute hierarchy amongst distinct behaviors, analogous to Maslow’s Hierarchy of Needs (Maslow, 1943). How would a fly prioritize when given multiple positive options? Would this choice be fixed or flexible? Much popular discussion on the disadvantages of multitasking and rapid task switching led us to examine whether flies finish one task before starting another or alternate between activities that satisfy competing drives. We also sought to determine if different decisions share common neural circuits or are controlled independently. We hypothesized that if decision-making itself, rather than performance of a specific behavior, were impaired, flies could show altered choices in multiple behavioral assays. Previous research examined decisions between two behaviors such as courting and feeding showed roles for specific tyraminergic neurons (Cheriyamkunnel et al., 2021) and gut hormones (Lin et al., 2022), but whether these same neurons have a more universal effect across multiple decisions was not known.

We designed behavioral choice assays to test a fly’s choices amongst grooming, feeding, and courting (**Fig. 1A-B**). All three of these behaviors are natural, productive actions, and the drive can be adjusted to perform each one by manipulating internal and external cues. We measured behavior onset, bout durations, and relative time spent grooming, feeding, or courting in pairwise assays. We proposed competing models to describe aspects of the decision-making process in fruit flies. These models present alternative possible answers to three questions: 1) Do fruit flies prioritize certain behaviors (**Fig. 1C**)? 2) Do different decisions share the same mechanisms (**Fig. 1D**)? 3) How do flies allocate time between two productive actions (**Fig. 1E**)?

**Figure 1:**
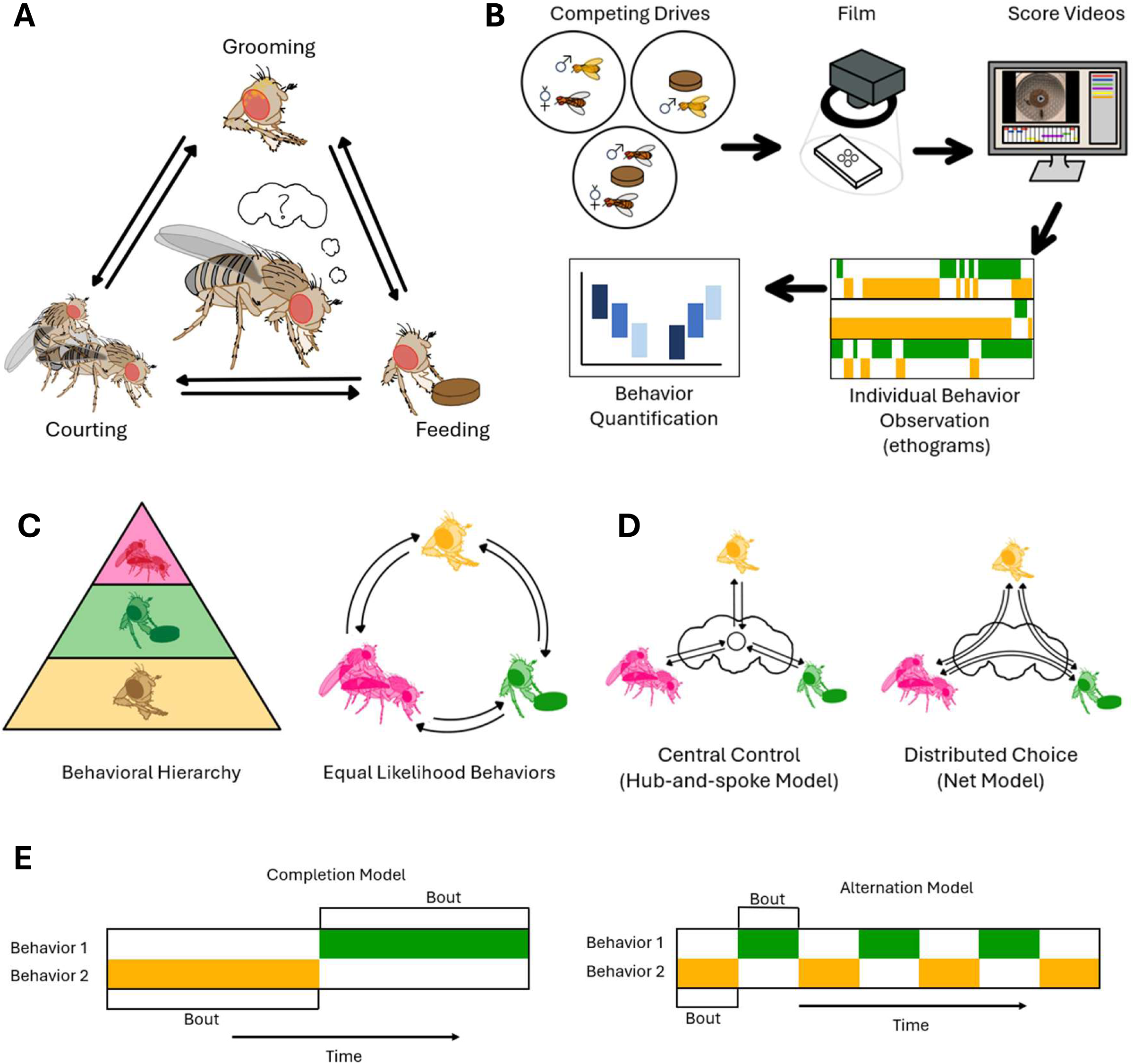
Conceptual overview and choice assay measuring how flies select among competing positive actions. A) A graphical abstract of the decision-making assay. B) An outline of the assay and analysis pipeline. C) Two models proposing the presence or absence of a behavioral hierarchy. D) Two models representing the decision-making process at an anatomical level: central vs distributed decision-making circuits. E) Two models hypothesizing about temporal distribution of behaviors resulting from competing drives: alternation vs sequential completion.

## Methods and Materials

### Fly strains

To study the behavior of wild type flies during decision making tasks, we used the *CantonS Drosophila melanogaster* strain (**Table 1**). The flies were raised on a mixture of cornmeal, molasses, yeast, and agar in a 25°C incubator on a 12-hour light/dark cycle. Within 24 hours of becoming adults, the males and females were separated into sex specific housing. Ice was used to anesthetize flies during sorting. All lines crossed to *UAS-CsChrimson* were kept in a dark 25°C incubator and raised on food containing 0.4 mM all-trans-retinal and starved on agar containing 0.4 mM all-trans-retinal. All lines crossed to *UAS-Kir2.1; TubP-GAL80^ts^* were kept at 30°C for 24 hours and then rested at room temperature (23°C) for 1 hour prior to filming.

**Table 1:** Fly Strains.

| **Table 1: Fly Strains** |  |  |  |
| --- | --- | --- | --- |
| **Strain** | **Nickname** | **Bloomington Stock ID** | **Obtained From** |
| CantonS | CantonS |  | Simpson Lab |
| pBDPGal4U in attP2 | Empty-GAL4 | 68384 | Simpson Lab |
| w[*] TI{RFP[3xP3.cUa]=2A-GAL4}Tbh[2A-GAL4.KI]/FM7a | Oct-GAL4 | 86145 | Bloomington |
| w[*]; P{w[+mC] = Tdc2-GAL4.C}2 | Tdc2-GAL4 | 9313 | Bloomington |
| w[1118]; TI{w[+mE.hs]=GAL4}TyrR[Gal4]) | TyrR-GAL4 | 67129 | Montell Lab |
| Th-GAL4 | Th-GAL4 |  | Simpson Lab |
| Trh-GAL4 (III) | Trh-GAL4 |  | Simpson Lab |
| w[1118]; P{w[+mC]=Lk-GAL4.TH}2M | LK-GAL4 | 51993 | Bloomington |
| w[1118]; P{y[+t7.7] w[+mC]=GMR65C07-GAL4}attP2 | LKR-GAL4 | 39344 | Bloomington |
| UAS-Kir2.1; Tub-GAL80ts | UAS-Kir2.1 |  | Louis Lab |
| w[1118]; P{y[+t7.7] w[+mC]=20XUAS-IVS-CsChrimson.mVenus}attP40 | UAS-CsChrimson | 55135 | Simpson Lab |
| w[*]; P{w[+mW.hs]=GawB}OK107 ey[OK107]/In(4)ci[D], ci[D] | OK107-GAL4 | 854 | Bloomington |
| pan[ciD] sv[spa-pol] |  |  |  |
| 201Y-GAL4 | 201Y-GAL4 |  | Simpson Lab |
| w[1118]; Pbac{WH}FoxPf03746 | FoxP | 85667 | Bloomington |

### Behavioral experiments

#### Filming conditions

For all assays, filming was done at room temperature (23°C). All filming was done with a FLIR Blackfly S USB3 camera and recorded top down with an infrared backlight at 30 fps.

Each fly was only used once in one assay. Flies were anesthetized by placing vials of flies on ice for transferring to either dusting chambers (grooming experiments only) or recording chambers, described in Seeds et al., 2014. All flies recovered from anesthetization for 15 minutes prior to filming to ensure maximum alertness. The same recording chambers were used for every experiment. All flies, male and virgin female, were 3 days old at the onset of filming and at the same subjective time of day to control for circadian effects.

#### Dusting

As described in the Seeds et al., 2014 paper, the dusting chambers consist of two 3D printed plates with 24 cylindrical wells each with a 15.6 mm diameter. Between the two plates lies a Nitex, 630 μm mesh sheet allowing for dust, light, and air to travel through the wells, but does contain the fly. The wells are sealed using aluminum sliders slid through grooves on the top plate. Anesthetized flies were placed in the four center wells, one fly per well. To dust the flies, 5 mg of Reactive Yellow 86 dust were added to the chambers containing alert flies. The chamber was then shaken in a reproducible manner to allow for even coating of dust on the flies. Once shaken, excess dust was removed by tapping the dusting chamber to allow dust to fall through the mesh. A corresponding filming chamber was then aligned with the dusting chamber to allow for transfer of the dusted flies. All dusting was performed under a Misonix WS-6 downflow hood.

The filming chambers used in this experiment are made of clear fabricated plates with four wells with a 15.6 mm diameter aligning with the four center wells of the dusting chamber. The chambers consist of three plates with a Nitex mesh sheet between the bottom two layers. The middle plate served as the chamber the fly resided in while the top plate and a clear slider acted as a lid which contained the fly but could be filmed through.

As soon as flies were transferred from the dusting chamber to the filming chamber, the filming chamber was then placed under the camera and filming commenced.

#### Grooming vs feeding

Prior to filming, flies were starved on agar (“wet-starved”: deprived of food but not water). *OK107-GAL4>UAS-Kir2.1; TubPGAL80ts*,*201Y>UAS-Kir2.1; TubPGAL80ts*, and their control strain cross progeny were starved for 12 hours, all other strains were starved for 24 hours. This time length was chosen based on a starvation titration performed on *CantonS* flies (**Fig. S1**).

Male flies were anesthetized and then placed in the dusting chambers and dusted using the dusting protocol described above. The dusted males were transferred to a filming chamber. In each well of the filming chamber, a small plug of cornmeal food was placed on the center of the mesh before the flies were introduced. The plug was made by inserting the wide end of a glass Pasteur pipette (7 mm diameter) into a standard food vial and then slicing the tube into disks approximately 2 mm thick. Fresh food was used for each experiment. Filming commenced as soon as flies were transferred.

#### Grooming vs courting

Both male and virgin females were anesthetized using ice. Males were transferred into a dusting chamber while females were transferred into a filming chamber. After allowing for 15 minutes recovery, the male flies were dusted using the above-described protocol. The female flies were not dusted because we found that the male flies did not initiate courtship to dusted females. Whether this is due to masked olfactory or pheromonal cues, altered visual appearance, or other factors is unclear. The male flies were then transferred directly into the chambers containing the virgin females; one male and one virgin female per well. Once flies were transferred, the filming chambers were placed under the camera and filming began.

#### Courting vs feeding

Only male flies were starved prior to filming. After being anesthetized, male flies rested in an empty filming chamber while females rested in a filming chamber containing a food plug as described above. Male flies were then shaken to simulate the shaking done in the grooming experiments. The male flies were then directly transferred into the chambers containing the female and the food. After transfer, the chamber was placed under the camera and filming began.

#### Triple competition assay

The protocol in this experiment was the same as described in the courting vs feeding protocol, except the males were dusted as in the dusting protocol prior to being transferred to the filming chamber consisting of a virgin female and a food plug. Filming began once flies were transferred.

### Screen

To study what types of neurons were involved in decision-making, a targeted screen was run using subsets found during a literature hunt (Pooryasin and Fiala, 2015; Cheriyamkunnel et al., 2021; Huser et al., 2012; Certel et al., 2010; Münch et al., 2022; Meschi et al., 2024). The GAL4/UAS system was used for both inactivation and activation of subsets of neurons. GAL4 males (Oct, Tdc2, TyrR, Th, Trh, LK, LKR, and Empty {Control}) were crossed with *UAS-Kir2.1; TubPGAL80ts* (inactivation) and *UAS-CsChrimson* (activation) virgin females. The resulting flies were then tested in all three decision-making assays. All strains in the screen had a *w1118* background.

#### Optogenetic activation

Prior to the optogenetic activation experiments with *UAS-Chrimson*, flies were raised on 0.4 mM all-trans-retinal. Within 24 hours of becoming adults, males were transferred to fresh tubes containing their regular cornmeal food mixed with all-trans-retinal while virgin females were transferred to a plain cornmeal food tube as described previously. During starvation experiments, male flies were transferred to agar tubes containing 0.4 mM all-trans-retinal mixed into the agar. During the experiment, an overhead FLDR-i132LA3 ring light (626 nm) producing approximately 0.85 mW/cm^2^ of red light was used for optogenetics activation. Prior to filming, flies were raised in the dark. During filming, the red light was on consistently throughout process.

### Data analysis

#### Video annotation

All videos were annotated for behavior manually using the VCode system (http://social.cs.uiuc.edu/projects/vcode.html) for the first five minutes. Courting, feeding, and grooming behaviors were annotated separately (i.e., the video was watched three times). Courting behavior was annotated for orienting (when a male stood facing a female within two body lengths), chasing (when a male walked towards a female within two body lengths), wing singing (when the male extends its wing to the side while facing the female), attempted copulation, and copulation. Feeding behavior was annotated for feeding (proboscis extension, contact, and retraction) and not feeding. Grooming behavior was annotated for anterior grooming (head sweeps and front leg rubbing) and posterior grooming (back leg rubbing, abdominal sweeps, wing cleaning, and thoracic sweeps); walking and standing behavior was also noted. Manual annotation was performed at a frame-by-frame resolution.

#### Ethograms

Ethograms showing the temporal distribution of behavior were made using MATLAB. Grooming, feeding, and courting are displayed on separate rows grouped by the individual fly to show instances of simultaneous behavior and to highlight the alternation pattern. Ethograms omitted walking behavior and showed all behavior subsets as their composite behavior courting, feeding, or grooming. Further analysis was done to look at both comparative amounts of each behavior as well as total behaviors. This statistical analysis and the creation of dot plots, box plots, Kaplan-Meier curves, preference indexes, and ternary plots were made using Excel.

#### Behavior initiation

Initiation was calculated as the first instance of a particular behavior. Wing singing, a courtship specific behavior, was used as the marker for courtship initiation because chasing and orienting behaviors can both be incidental of two free moving flies in a small space. Feeding initiation was marked as the first time a male fly’s proboscis contacted food. Grooming initiation was not analyzed because eye cleaning commences instantly and could not be captured before filming began. All flies used in **Figure 5** were *CantonS* flies. Behavior initiation was shown using a Kaplan-Meier curve. This graph, though most used in the medical field, was selected because it tracks time to incidence while also accounting for lack of incidence, i.e. when the fly never fed or wing sang.

#### Bout length analysis

Bout lengths were calculated as time a fly continuously performed an action. Any standing or walking breaks in behavior for less than a second were disregarded. Copulation was not included in courtship bout length analysis. Outliers were excluded and how many and when are noted in each figure legend. Outliers were defined as bouts that were above the upper quartile. No more than two bouts per experiment were excluded. All flies used in **Supplemental Figure 2** were *CantonS* flies.

#### Preference index

The preference index tracks which of two behaviors in the pairwise competition assays was preferred. This calculation was done by (Behavior A - Behavior B)/(Behavior A + Behavior B). A preference index of 1 indicates the fly performed Behavior A but never Behavior B, an index of −1 indicates the fly performed Behavior B but never Behavior A, and an index of 0 indicates the fly spent equal time on each behavior.

#### Ternary plot

Though most often used in physical sciences, we chose this plot in **Figure 8B** because it portrays a three-dimensional preference index for the triple competition assay. Each data point has three coordinates adding up to 100. Each coordinate indicates the percentage of time spent on a behavior of the time spent on any of the three behaviors. The coordinates are determined by calculating [Behavior A – (Behavior B + Behavior C)]/(Behavior A + Behavior B + Behavior C) x 100 for each of the three behaviors for each fly.

#### Statistical analysis

P values were calculated for two groups with the Mann-Whitney test. For more than two groups, a Kruskal-Wallis test followed by a Dunn’s multiple comparison test and Bonferroni correction were performed. The p value for the Kaplan-Meier curve was found using a log-rank analysis followed by a Bonferroni correction. The dot plots in **Figures 2B**, **3B**, and **4B** serve to demonstrate variation in the population, and because each dot conveyed time spent for two different behaviors, p values were not found for these plots.

**Figure 2:**
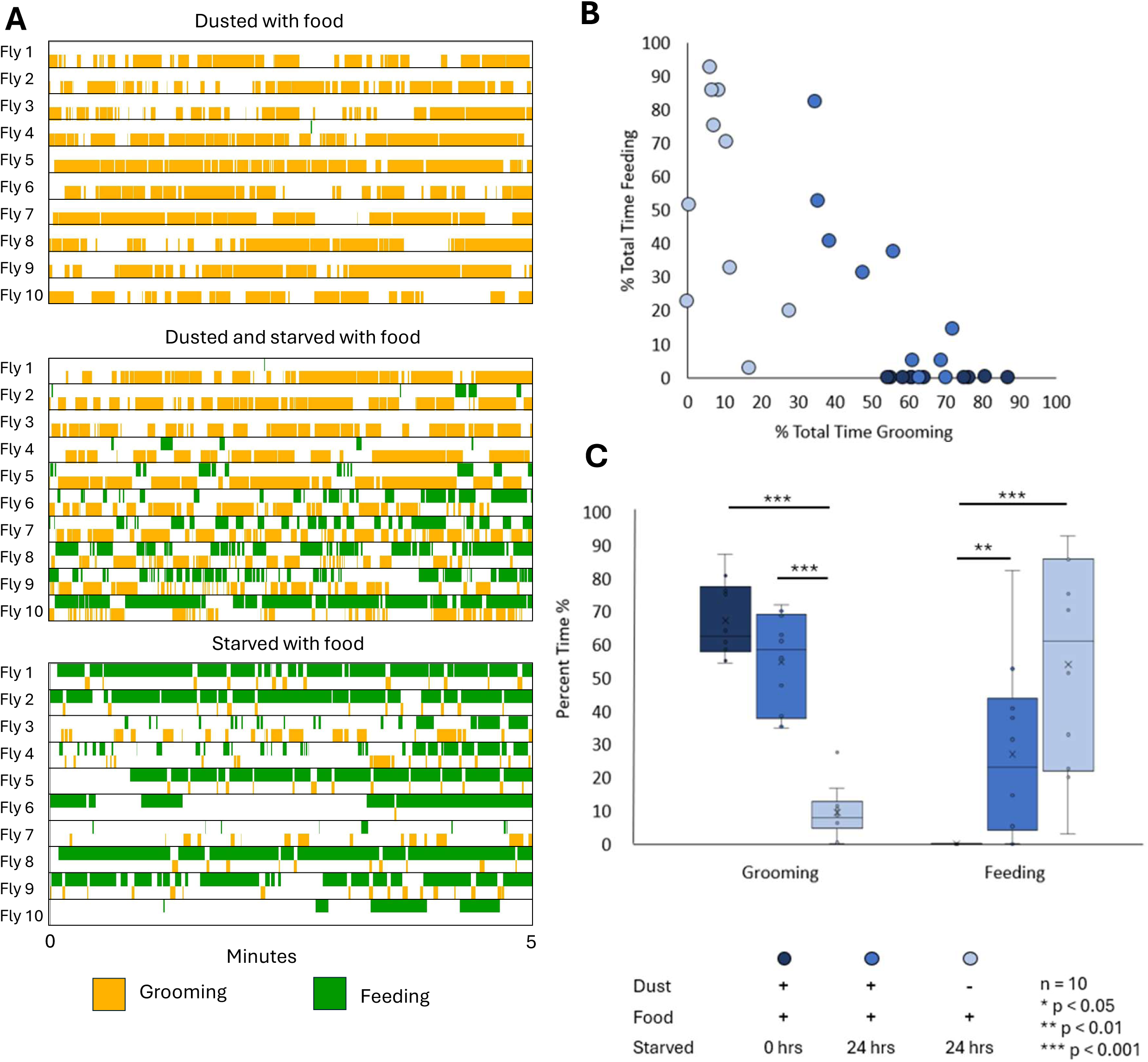
Grooming and feeding behaviors alternate when drives compete. A) Ethograms depicting the first five minutes of the grooming vs feeding assay. Male flies are provided with food and are starved or dusted as indicated. B) Dot plot comparing the percent of total time spent grooming to the percent of total time spent feeding for each individual fly. C) Box plot comparing the average time spend for all flies. X denotes mean, bar denotes median, and whiskers denote the lower and upper quartile. Statistical significance was assessed using a Kruskal-Wallis test followed by a Dunn’s multiple comparison test and Bonferroni correction.

## Results

### Characterization of pairwise behavioral choices

We first established what flies do in our assay chambers when only a single behavioral option is presented (**Fig. 2A**). When flies are covered in dust or debris, and not starved, they groom. There is individual variability in timing and amounts (**Fig. 2B**), but the overall grooming behavioral response is robust: most flies spend 60-80% (**Fig. 2C**) of their time grooming, often in long bouts in a 5-minute assay. We hypothesize that the mechanosensory stimulus from the dust provides strong external drive.

When flies are starved but not dusted, hunger is a strong internal drive and flies spend time feeding to satisfy it (**Fig. 2A**). There is individual variability (**Fig. 2B**) but most flies spend 20-80% of their time feeding (**Fig. 2C**), often in long bouts, in a 5-minute assay.

We hypothesize that variability is caused by differences in fat storage or exact time of last meal before experimental food deprivation began.

We then examined how flies choose between grooming and feeding when both drives are present. When both dusted and starved, the ethograms, records of the actions the flies perform over the assayed time (**Fig. 2A**), show that the flies alternate between bouts of grooming and feeding, supporting the Alternation Model proposed in **Figure 1E**. The ratio of behaviors for each individual fly is variable, as shown in **Fig 2B**, but on average, most flies spend 40-70% of their time grooming and 5-45% of their time feeding (**Fig. 2C**), often in shorter alternating bouts between feeding and grooming (**Fig. S2**). Therefore, the total time spent grooming and the total time spent feeding are both reduced when compared to performance without competition. We conclude that the presence of two competing drives causes the flies to alternate, dividing their time between behaviors to reduce both motivating drives.

Similar results occur when grooming and courting drives compete (**Fig. 3**). Males alternate between grooming and courting (**Fig. 3A**). Individual flies demonstrate variability in the courting behavior (**Fig. 3B**), which we hypothesize is due in part to the response of free roaming virgin females. On average, clean male flies spend 90% of their time courting (**Fig. 3C**) when they share a small chamber with a conspecific virgin female, but when the male is covered with dust, courting is reduced, and grooming is increased, suggesting a trade-off in time spent on each behavior.

**Figure 3:**
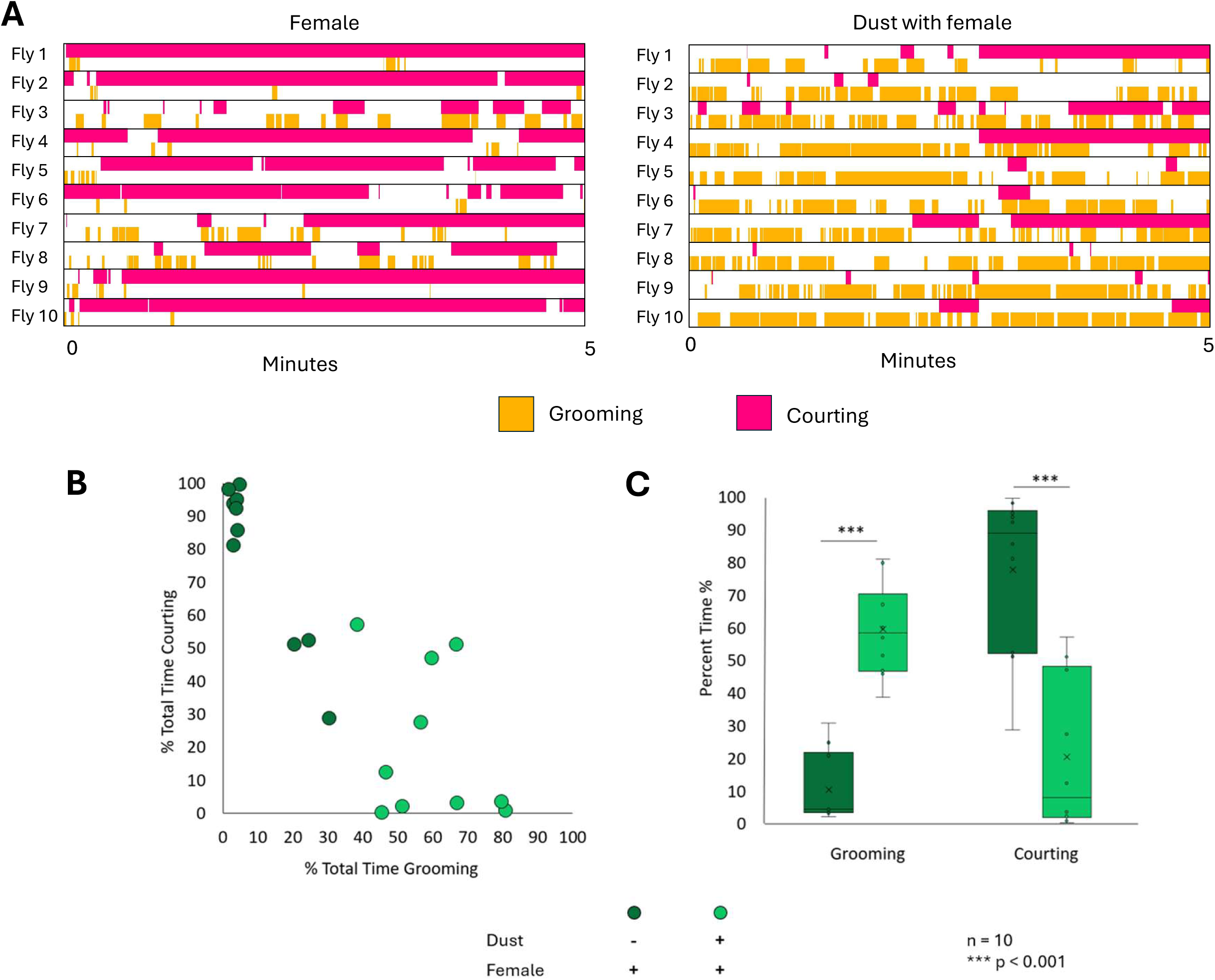
Grooming and courting behaviors alternate when drives compete. A) Ethograms depicting the first five minutes of the grooming vs courting assay. Male flies were raised in sex isolation and presented with a virgin female. Dust is present when indicated. B) Dot plot comparing percent of total time spent grooming to the percent of total time spent courting for each fly. C) Box plot comparing percent of total time spent on each behavior. X denotes mean, bar denotes median, and whiskers denote the lower and upper quartile. Statistical significance was assessed using a Mann-Whitney test.

We also see alternation in the courting vs feeding assay. Starved male flies will alternate between courtship and feeding when given access to both virgin females and food (**Fig. 4A**). Most males alternate between courting and feeding with bouts of shorter duration (**Fig. S2**), rather than feeding for a sustained period before switching to courtship. As in the grooming vs feeding assay, the amount of time spent feeding is variable (**Fig. 4B**). The total amount of time spent courting is more significantly reduced when food is available (**Fig. 4C**).

**Figure 4:**
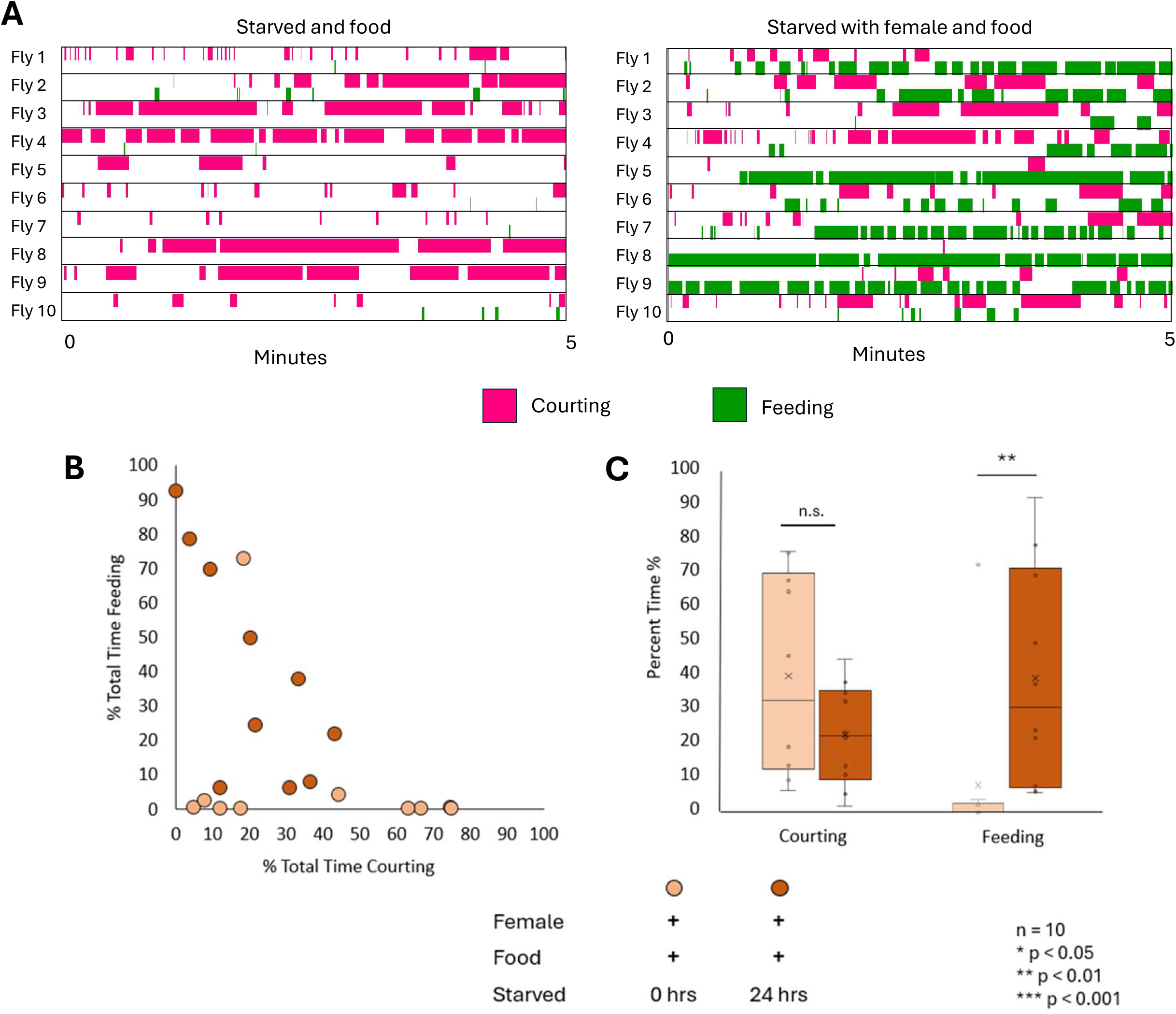
Courting and feeding behaviors alternate when drives compete. A) Ethograms depicting the first five minutes of the courting vs feeding assay. Male flies have been starved and raised in sex isolation and have been presented with a virgin female and food. B) Dot plot comparing the percent of total time spent courting to the percent of total time spent feeding for each fly. C) Box plot comparing percent of total time spent on each behavior. X denotes mean, bar denotes median, and whiskers denote the lower and upper quartile. Statistical significance was assessed using a Mann-Whitney test.

During the three pairwise competition assays, we observe rare attempts to multitask: occasionally a fly will perform posterior grooming during feeding or courtship singing and occasionally feed while courtship singing. These combinations are physically possible (**Fig. S3**) but infrequent.

### Behavior initiation is delayed by competing drives

We next evaluated flies’ response to competition using behavior initiation time. We measured when they initiate courtship (**Fig. 5A**) or feeding (**Fig. 5B**) with Kaplan-Meier curves to depict percentage of flies that have initiated the behavior at each timepoint. (Note that we used wing-song to mark the onset of courtship because it is more specific to courtship than orienting or following, and we could not measure grooming onset time accurately because it always began as soon as the flies were exposed to dust and before we could begin filming).

**Figure 5:**
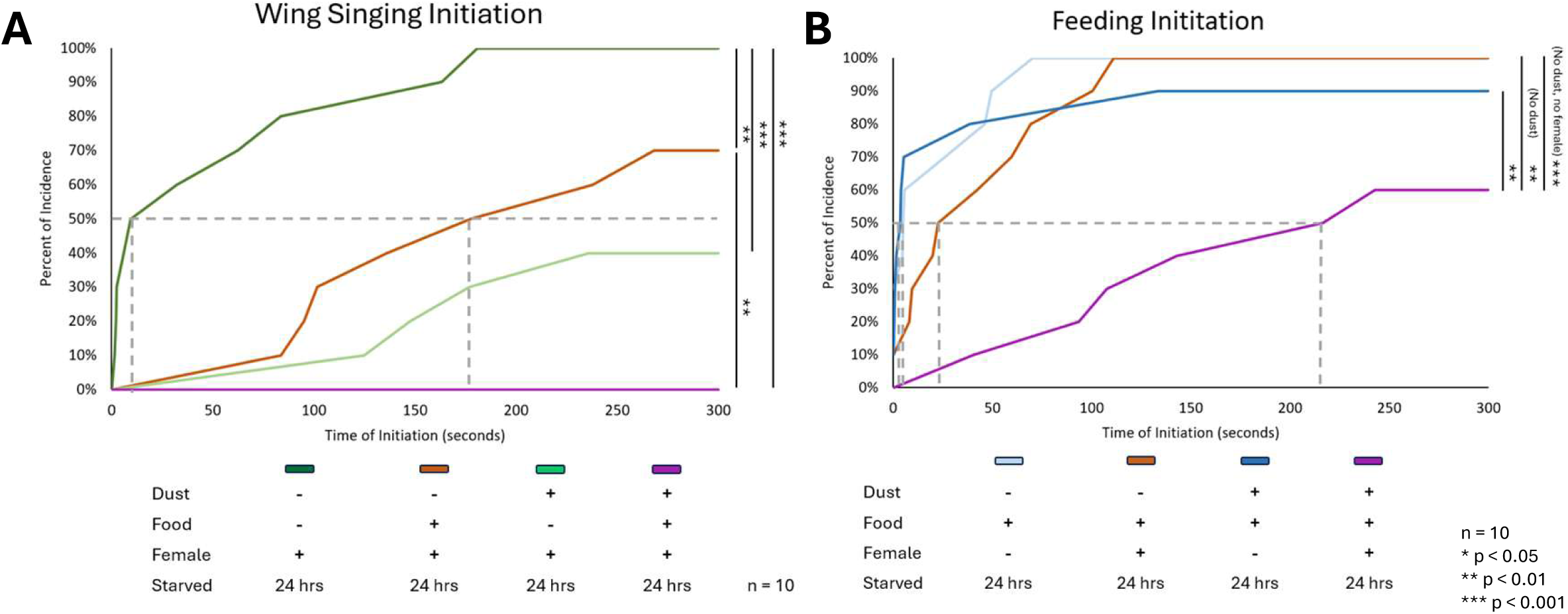
Competing drives result in delayed behavior initiation. A) A line graph depicting a Kaplan-Meier curve tracking initiation of wing singing courtship behavior. The dotted line represents the 50% mark. B) A line graph depicting a Kaplan-Meier curve tracking initiation of feeding behavior. The dotted line represents the 50% mark. Statistical significance was assessed using a log rank analysis and a Bonferroni correction.

In all of our assays, flies predominantly initiated grooming first, followed by feeding, and then courting. This order could reflect priority or opportunity: the flies are covered in dust, so they receive the mechanosensory stimulation that drives the grooming response immediately. Since the chambers are small (15.6mm in diameter), males typically encounter both the moving female and the stationary food patch quickly. Therefore, we believe the initiation and allocation of time spent is primarily due to relative drive strength or priority rather than just proximity.

The average time of courtship initiation is delayed by the presence of food or dust (**Fig. 5A**), while the initiation time of feeding is delayed when more than one alternative is present: dust and virgin females (**Fig. 5B**). In a triple competition assay (where the flies were starved, dusted, and presented with females and food), the flies never initiated wing singing (**Fig. 5A**) and delayed feeding (**Fig. 5B**): only 60% of flies fed at all. Thus, measurements of change in behavior initiation time support the analysis of total time spent in each behavior to indicate that flies faced with competing drives alternate between the actions required to satisfy them as proposed in **Fig. 1E**.

Bout length is another temporal feature that shows the effects of competition. The amount of time a fly spends performing each behavior is reduced as competition increases (**Fig. S2**). Feeding and courting bouts become shorter. For example, feeding bouts average 13.6 s without competition, 10.4 s when a female is present, 6.30 s when food was present, and 3.55 s in triple competition. Average courtship bout length is reduced from 4.82 s to 1.19 s when dust was present or 1.74 s when food was present and 0.59 s when both dust and food are available. Grooming bouts are also reduced by competition, but less so (from 12.45 s to 9.05 s with female, to 7.27 s with food, and to 9.54 s with both). Both onset time and bout length stability support a soft hierarchy of grooming, then feeding, then courting as proposed in **Figure 1C**.

### Candidate genes, neurons, and brain regions may affect aspects of decision-making

#### Candidate genes

To further study the decision-making behavior of *Drosophila melanogaster*, we considered candidate genes, neurons, and brain regions. Previous experiments implicated mutations in the *FoxP* transcription factor in the time course of perceptual decisions when sensory stimuli were ambiguous (Palazzo et al., 2020). We tested whether available *FoxP* mutations behaved differently in our pairwise competition assays. The genetic background of the mutants affected the starvation conditions required to balance the choice between feeding and grooming or courting, but the primary effect we observed was an increase in the average bout duration of anterior grooming, perhaps indicating that transitions between anterior and posterior grooming actions was impaired (**Fig. S4**). This observation encouraged us to investigate bout duration as another manifestation of decision-making, potentially acting over a shorter timescale and within an overarching behavioral context (**Fig. S4**). *FoxP* mutants do not show differences in action selection within courtship, indicating the grooming effect is not due to a wider switching impairment (*data not shown*). Our initial grooming bout duration results were observed with a hypomorphic *FoxP* allele, f03746 (Mendoza et al., 2014), and when we were unable to confirm them with the additional knock-out allele 5-SZ-3955 (DasGupta et al., 2014), we did not pursue this gene candidate further.

#### Candidate neurons

A fortuitous observation in an undergraduate lab class showed that activation of the neurons targeted by *leucokinin-GAL4* caused reduced grooming in response to dust – and an increase in proboscis extension. This led us to test whether these neurons affected the choice between feeding and grooming. We conducted a small screen of candidate neuron subsets including neurons expressing leucokinin and leucokinin receptor, tyramine and tyramine receptor (Cheriyamkunnel et al., 2021), and other aminergic neurons (Gautham et al., 2024) (**Fig. S5**). We silenced and activated these neuron populations in the three pairwise competition assays, corroborating the published findings on tyraminergic neurons in a new assay and quantifying the anecdotal observations on the neurons expressing *leucokinin-GAL4*.

*Leucokinin-GAL4* drives expression in more than 20 neurons in the brain and ventral nerve cord (Nässel, 2021). Leucokinins are a group of neuropeptides associated with feeding – satiation and termination, sleep, nociception, and memory formation. Leucokinin has been implicated in the interaction of hunger and noxious heat (Ohashi & Sakai, 2018) and in sexual receptivity in juvenile females (Chen et al., 2025).

In single behavior assays, silencing Leucokinin neurons (*leucokinin-GAL4*>*UAS-Kir2.1;tubGAL80ts)* results in a decrease in feeding behavior (**Fig. 6A**) while acute activation (*leucokinin-GAL4>UAS-CsChrimson)* increased feeding and reduced grooming (**Fig. 6B**). Activation of the neurons expressing *leucokinin-GAL4* produced flies that had no noticeable motor deficits in grooming, courting, or feeding assays, but continuously extended their proboscis parallel to the surface.

**Figure 6:**
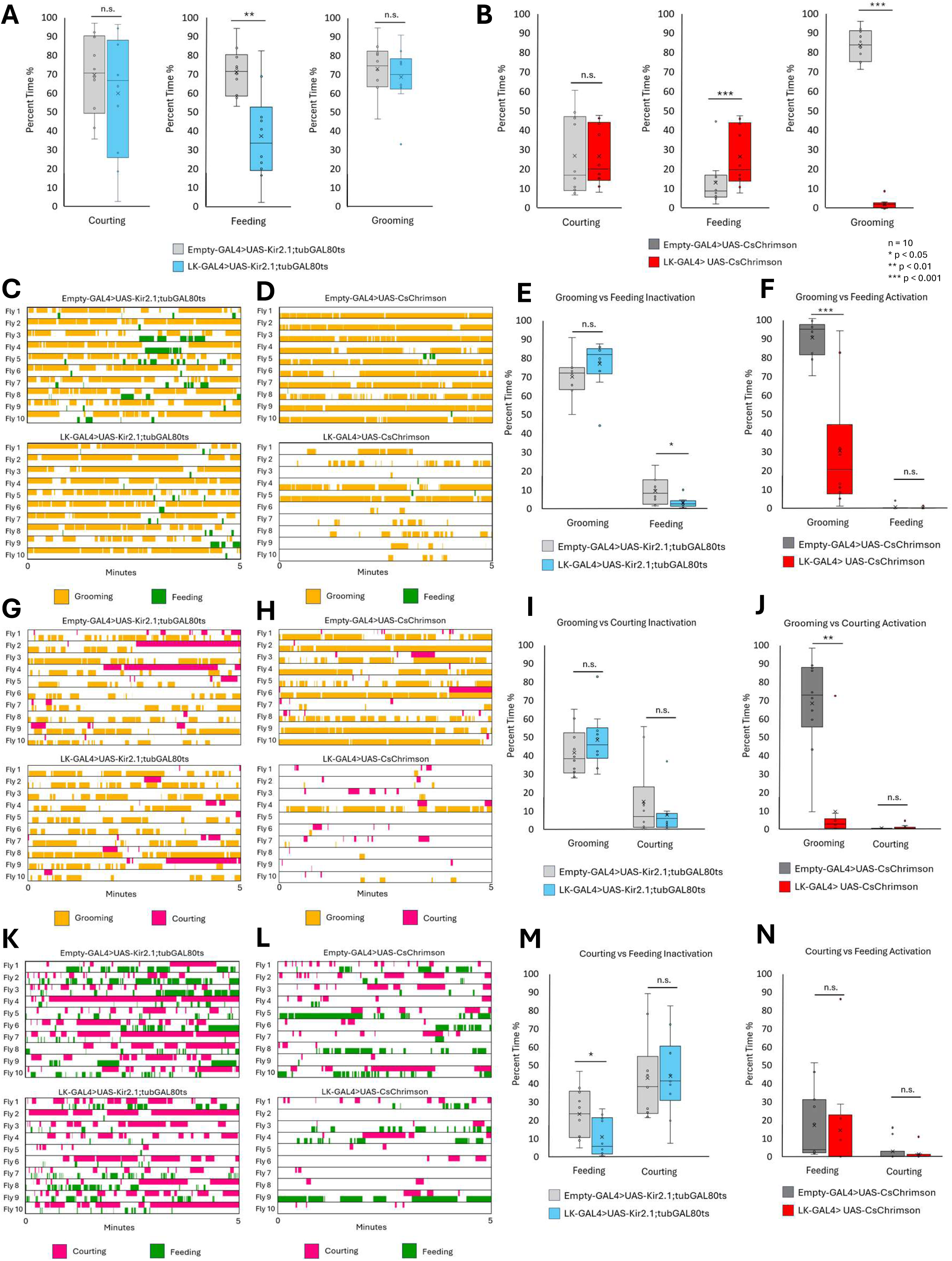
Inactivation and activation of leucokinin producing neurons in the three choice assays. A) Single drive analysis of inactivated leucokinin producing neurons. B) Single drive analysis of activated leucokinin producing neurons. C) Grooming vs feeding inactivation ethograms. D) Grooming vs feeding activation ethograms. E) Box plot corresponding to C. F) Box plot corresponding to D. G) Grooming vs courting inactivation ethograms. H) Grooming vs courting activation ethograms. I) Box plot corresponding to G. J) Box plot corresponding to H. K) Courting vs feeding inactivation ethograms. L) Courting vs feeding activation ethograms. M) Box plot corresponding to K. N) Box plot corresponding to L. In the box plots, the X denotes mean, bar denotes median, and whiskers denote the lower and upper quartile. Statistical significance was assessed using a Mann-Whitney test. All figures are n = 10.

In pairwise competitions, we observed trends consistent with the single behavior assays: silencing caused decreases in feeding (**Fig. 6C, E, K, M**) while activation decreased grooming (**Fig. 6D, F, H, J**). While these neural manipulations could affect either behavior prioritization or execution, we favor execution because we did not observe reciprocal increases in competing behaviors. That the reduction of grooming is not accompanied by an increase in feeding (**Fig. 6F)** or courtship (**Fig. 6J**) could be due to a general failure to switch between behaviors or a motor deficit, we do not believe this explanation is likely because alternation was normal in the courting vs feeding assay (**Fig. 6L, N**).

Though there was no significant change observed in the percentage of total time spent on courtship in either the individual or competition assays, we observed a more subtle change in the courtship micro-behaviors (**Fig. S6**): more late-stage behaviors such as attempted copulation and copulation were observed when leucokinin neurons were silenced (**Fig. 6K, M**) and fewer late-stage behaviors were observed when leucokinin neurons were activated (**Fig. 6L, N**).

#### Candidate brain regions

The mushroom bodies are a brain area involved in associating positive and negative valence to olfactory cues in associative learning and memory paradigms, as well as a wealth of other higher order adaptive behaviors (Zhang et al., 2007; Parnas et al., 2024; Suárez-Grimalt et al., 2024; Saito et al., 2022; Kasoch et al., 2015; Chan et al., 2024). We chose to test this brain region as a candidate for a central control region for decision-making.

We tested whether silencing mushroom body neurons (using broad drivers *201Y*- and *OK107-GAL4*) would alter grooming, feeding, or courtship in individual behavior assays or competitive choice. We saw a strong reduction in percentage of total time spent courting in the individual assay (**Fig. 7A**) but not the grooming vs courting or courting vs feeding choice assays (**Fig. 7C, F, D, G**). The increase in feeding behavior (**Fig. 7A**) observed in the single behavior assay was not observed in the competition assays (**Fig. 7B, E, D, G**). These results suggest that the changes caused by mushroom body inactivation may be due to reduced locomotor activity rather than defects in decision-making or changes in priority.

**Figure 7:**
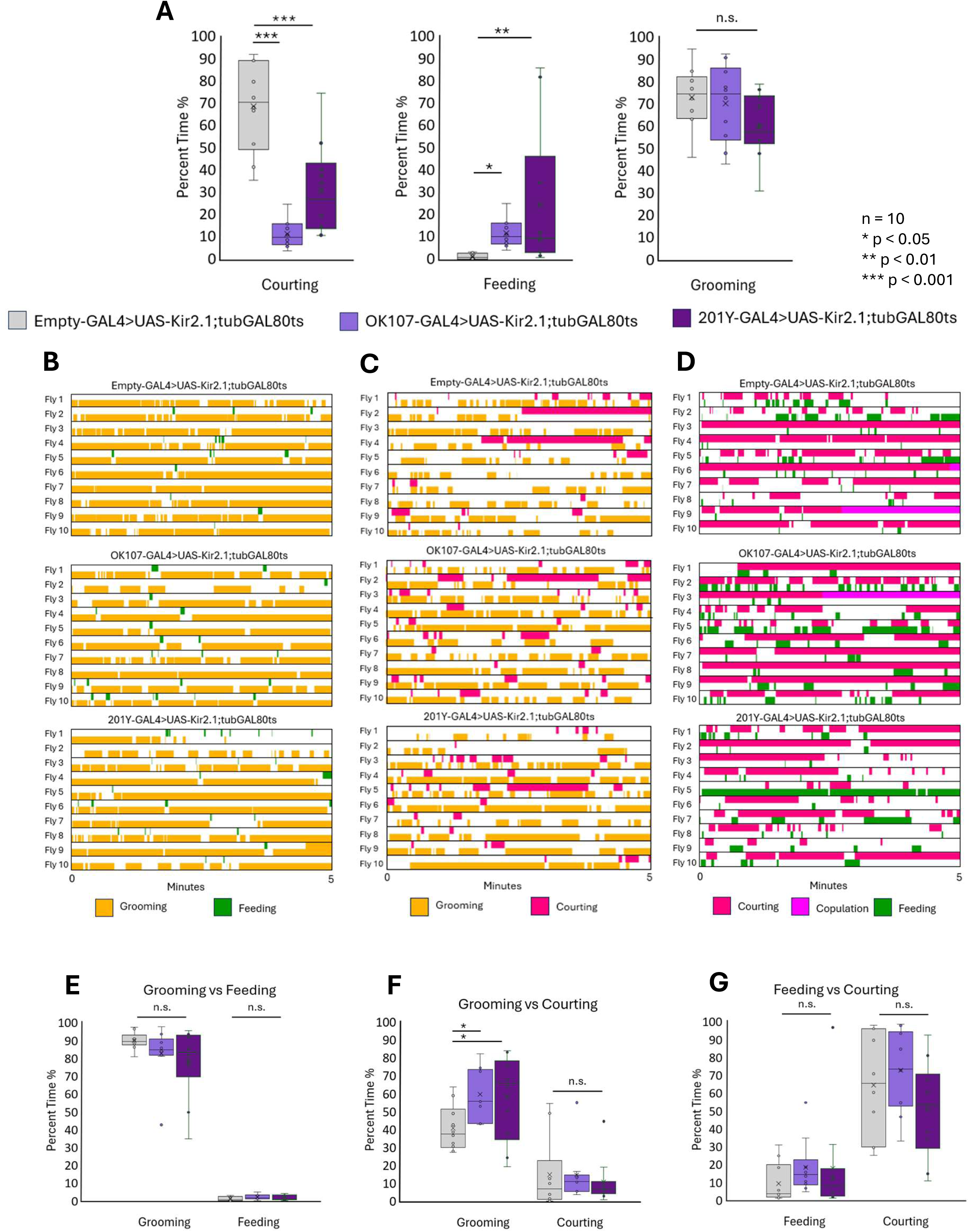
Inactivation of mushroom bodies in the three choice assays. A) Single drive analysis of inactivated mushroom bodies. B) Grooming vs feeding inactivation ethograms. C) Grooming vs courting inactivation ethograms. D) Courting vs feeding inactivation ethograms. E) Box plot corresponding to B. F) Box plot corresponding to C. G) Box plot corresponding to D. In the box plots, the X denotes mean, bar denotes median, and whiskers denote the lower and upper quartile. Statistical significance was assessed using a Mann-Whitney test. All figures are n = 10.

### The effect of genetic background on the triple competition assay

The strains used in the experiments in this paper have two different genetic backgrounds *CantonS* (**Fig. 2-5**) and *w1118* (**Fig. 6-7**). The *CantonS* flies were used to initially characterize the decision-making behavior in the pairwise competitions. The GAL4/UAS strains with a *w1118* background were used to target the effects of specific genes, neuron subsets, and brain regions on decision-making. Comparing the two sets of data we observed the *w1118* background flies were overwhelmingly grooming when dust was introduced (**Fig. 6-7**). We know this is not due to the flies not being starved enough or interested in the females because they performed well in the courting vs feeding assay as well as the individual behavior assays. We hypothesize that the *w1118* flies have a higher priority on grooming than the other behaviors, which would also support our conclusion for the *leucokinin-GAL4* results (**Fig. 6**).

To test the effect of the genetic background, we conducted a triple competition assay comparing *CantonS* flies and *w1118* flies (**Fig. 8A**). We observed that the *w1118* flies had a greater preference for grooming than the other two behaviors. To plot this, we used a ternary plot to show the ratio of time spent between the three behaviors (**Fig. 8B**). To read, follow the grey line headed towards 100 for each behavior. Resulting three coordinates will total 100. Ex. The *w1118* outlier sits at 45 courting, 28 feeding, and 27 grooming. We also observed that the percent of total time spent grooming increased for *w1118* and feeding decreased (**Fig. 8C**). Courting remained unchanged and the flies never performed any courting specific behaviors (orienting and chasing can be incidental). We also looked at bout lengths for each of the three behaviors. There was an increase in grooming bout durations in the *w1118* flies (**Fig. 8D**) and no change in the other two behaviors (**Fig. 8E-F**). The *w1118* grooming bout length is twice as long as the *CantonS* grooming bout length and longer than the *CantonS* grooming only behavior as well (**Fig. S2**). This coincides with the previous remarked upon data (**Fig. 2A, 3A, 4A, S2**) where we observed that as priority lowered (competition introduced) bout length lowered. We hypothesize, that in *w1118* we observe an increase in bout length because there is an increase in priority.

**Figure 8:**
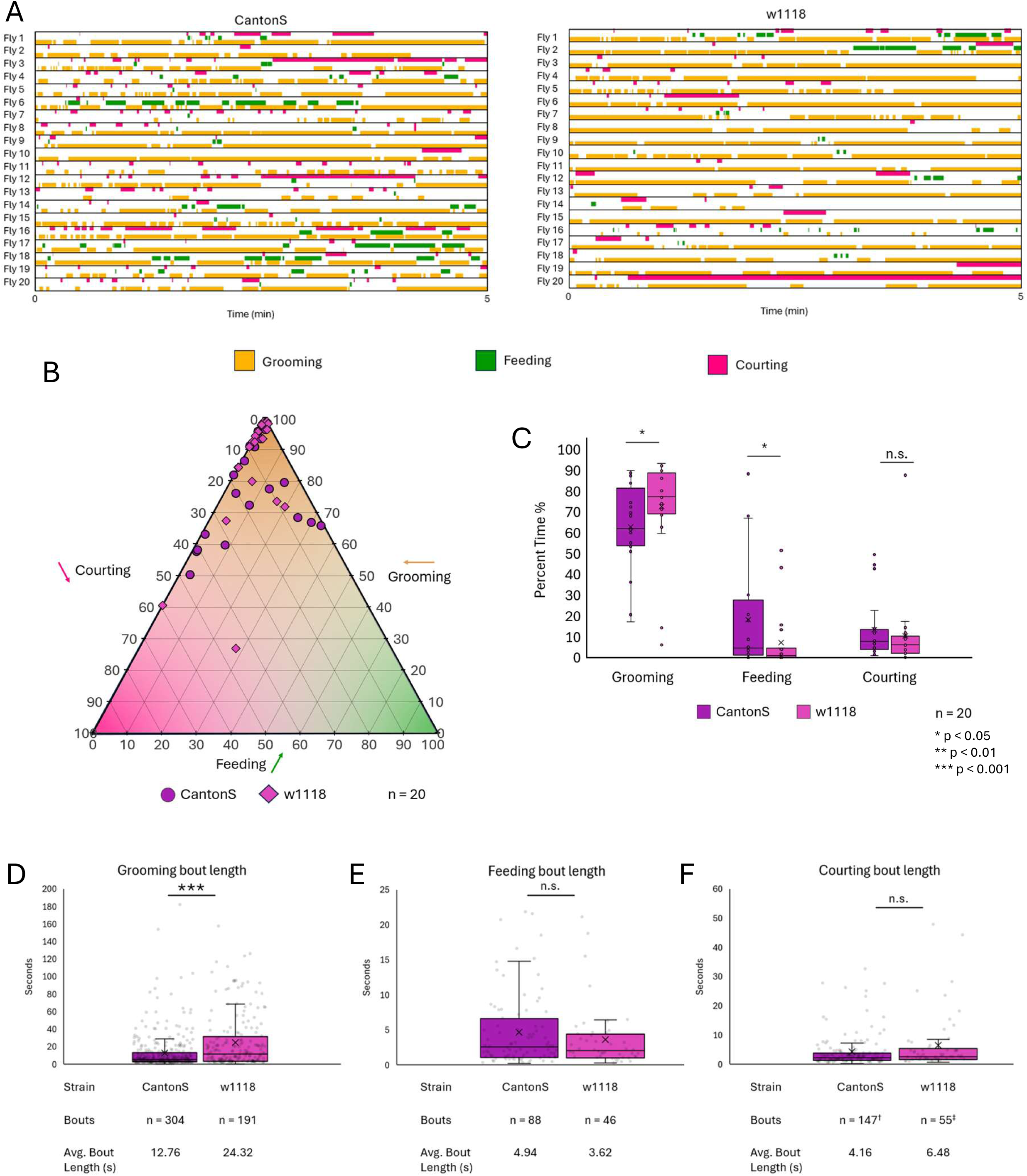
Effects of genetic background on the triple competition assay. A) Ethograms depicting *CantonS* and *w1118* flies in the grooming vs feeding vs courting triple competition assay. B) Ternary plot of individual *CantonS* and *w1118* flies depicting percent of time spent among three competing behaviors totaling 100%. C) Box plots comparing percent of total time spent on grooming, feeding, and courting during the first five minutes of the assay. D) Box plot with a dot plot overlay depicting the length of individual bouts of grooming during the triple competition assay. E) Box plot with a dot plot overlay depicting the length of individual bouts of feeding during the triple competition assay. F) Box plot with a dot plot overlay depicting the length of individual bouts of courting during the triple competition assay. ^†^2 outlying bouts were removed ^‡^1 outlying bout was removed. In the box plots, the X denotes mean, bar denotes median, and whiskers denote the lower and upper quartile. Statistical significance was assessed using a Mann-Whitney test.

## Discussion

### Characterization of pairwise decision-making

We characterized decision-making behavior in *Drosophila melanogaster* males during pairwise behavior competition assays. We described how the internal states of hunger and mating drive interact with external stimulation from dust in the presence of different environmental resources such as food and virgin females to influence decision-making. We show that flies alternate between behaviors (**Fig. 2A, 3A, 4A**): we propose that they do this as their actions satisfy competing drives. Other explanations are possible: the drives may decay or habituate with different time-courses, or arousal thresholds may affect each behavior differently. Competition resulted in changes in time allocation across behaviors (**Fig. 2C, 3C, 4C**) and delayed behavior initiation (**Fig. 5**). The behavior alternation indicates that while flies do have a hierarchy of needs, they can adapt their responses based on context (**Fig 1C, E**). This contextual flexibility resembles need-based prioritization of feeding, social behaviors, and defensive responses in mice (Burnett et al., 2013), indicating that this may be a general, adaptive mechanism across species. We find individual variability in behavior choices, even in genetically homogenous flies subjected to the same experimental conditions (**Fig. 2B, 3B, 4B**). Feeding is the most variable behavior, which we propose is due differences in hunger drive from timing of last meal before starvation period or fat storage capacity. We also report that flies can perform two behaviors simultaneously (**Fig. S3**), suggesting that these control circuits are not exclusive.

### Effects of genes and neurons on decision-making

We tested some candidate genetic mutations (*FoxP*) and neuronal populations (leucokinin, aminergic, and Mushroom Body neurons) for defects in decision-making. In the *FoxP* mutants, we saw no change in decision-making overall but observed longer bout durations of anterior grooming (**Fig. S4**), suggesting an avenue for further investigation. Manipulating activity in the neurons expressing the leucokinin neuropeptide affects performance of individual behaviors: silencing reduces feeding while activating reduces grooming (**Fig. 6A, B**); these effects on individual behavior performance likely account for the changes in the pairwise choice assays (**Fig 6E, F** and **J**). We do not yet know whether this reflect altered drive or ability to execute. We also observed that silencing leucokinin neurons accelerated onset of late-stage courtship behaviors (**Fig. S6**). These results suggest that leucokinin neurons warrant further investigation. We also tested whether activity of aminergic neurons affects decision-making in our assays (**Fig. S5**). Our results align with previously published work implicating tyraminergic neurons in choices between feeding and courtship. Because the mushroom body neurons have established roles in olfactory perceptual decision-making and learned association with valence, we tested inactivation of these neurons in our assays (**Fig. 7**). Single behavior analysis revealed a decrease in courting and an increase in feeding behaviors, but the competition assay between courting and feeding showed no significant changes. We hypothesize that mushroom body inactivation affects behavior performance rather than drive strength or decision-making, but confirmation requires further experiments.

We expected that disruption of a central decision-making center would affect choices in multiple assays and none of the neurons we tested produced this outcome. Therefore, we did not uncover any evidence identifying such a hub. But we did not rule out its existence, either. We tested a limited set of neuronal populations, so perhaps a broader screen would uncover neurons with a more global effect. Alternatively, the expectation of a global effect may be flawed: if decision-making is distributed or shows functional redundancy, neurons that affect only one of our assays may still contribute to decision-making more broadly. Our current results neither confirm nor exclude a central mechanism for decision-making (**Fig. 1D**).

### Genetic background affects decision-making

Our mutations and transgenic lines were in a different genetic background (*w1118*) from the *CantonS* wild-type flies we used to establish the initial choice assay conditions. We observed increased grooming in all of the assays with *w1118* flies relative to *CantonS*. Although flies with the *w1118* background were capable of all the individual behaviors, in the competition assays, grooming tended to dominate. To further elucidate effects of genetic background, we tested *w1118* and *CantonS* flies in a triple competition assay: grooming vs feeding vs courting (**Fig. 8**). We found that the *w1118* flies groomed more, fed less, and had longer bouts of grooming, indicating that genetic variants can influence behavioral priorities or performance. While we did not map genetic causes contributing to these differences, our observations underscore the importance of considering genetic background and comparing to appropriate controls.

## Conclusion

The neural basis of decision-making remains an open and challenging area of investigation, with new experiments representing different levels of analysis and employing the respective strengths of different animal models (International Brain Laboratory et al., 2025). Our experiments here further characterize decision-making behavior in *Drosophila melanogaster*. We show that fruit flies do have a hierarchy among the three behaviors tested but show flexibility in response to drive strengths and available resources. We show that flies alternate among competing behaviors. And we do not identify or exclude a central locus that affects decision-making. These assays and results guide future exploration into both behavioral algorithms and neural circuit mechanisms controlling how decisions are made.

## Acknowledgements

We would like to acknowledge our lab mates Neil Zhang, Sunanda Marella, Li Guo, Varun Sane, Shingo Yoshikawa, Primoz Ravbar, Durafshan Sayed, and Diego De Alba for their support and continuous feedback. We thank the UCSB undergraduates who helped score behavior video: Omar Alkhairi, Dipakshi Dhinakaran, Armaan Lebaschi, Taylor Pettway, Rose Rezaee, Rachel Tajiri, Kylie Viray, Jiaqi Wu, Sierra Gordon, Shuwei Lin, Meier Mai, Ryan Nishioka, Giulia Pellegrini, and Yifan Xu. We would also like to thank the Montell Lab and the Louis Lab for sharing their strains and Dr. Sung Soo Kim for his advice on statistical tests.

## Data and resource availability

Data and resource availability: All relevant data and details of resources can be found within the article and its supplementary information. Reagents available upon request.

