## Supplementary Data for "Behavior choices amongst grooming, feeding, and courting in *Drosophila* show contextual flexibility, not an absolute hierarchy of needs"

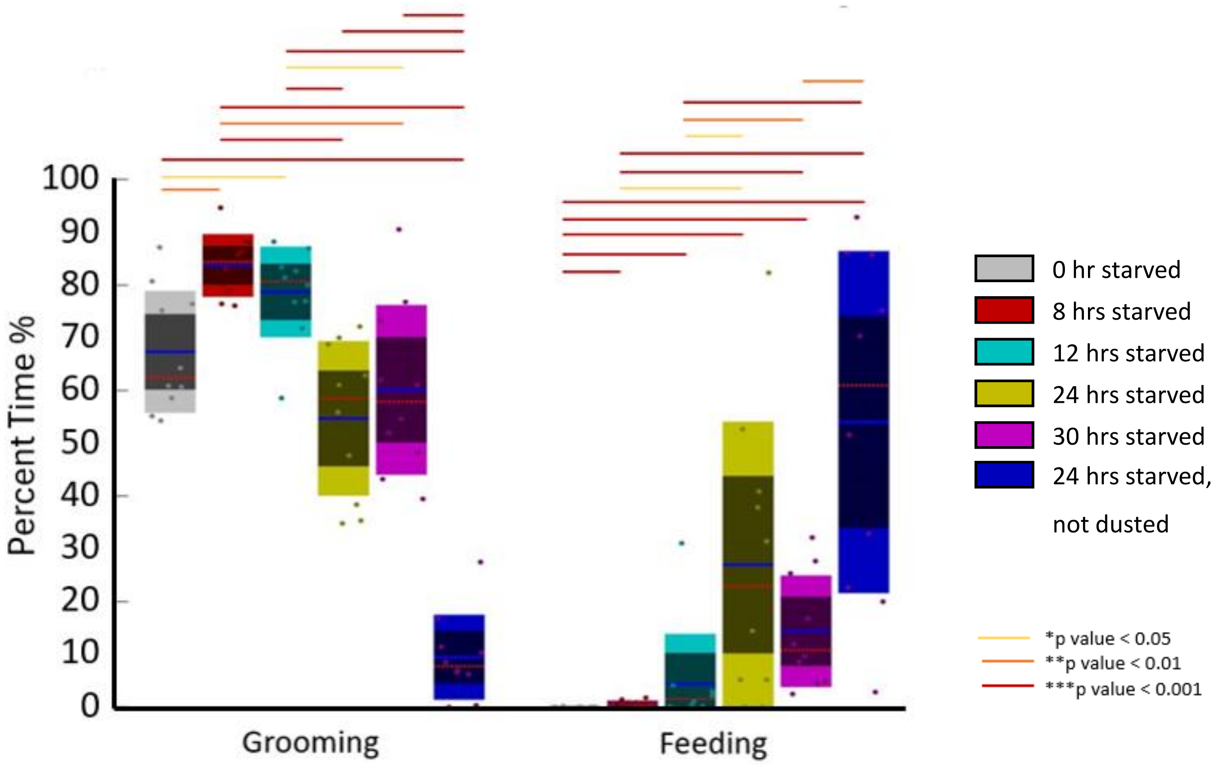

100
90
80
70
60
50
40
30
20
10
0
Percent Time %
Grooming
Feeding
0 hr starved
8 hrs starved
12 hrs starved
24 hrs starved
30 hrs starved
24 hrs starved, not dusted
*p value < 0.05
**p value < 0.01
***p value < 0.001

**Supplemental 1: CantonS behavior changes with respect to hunger drive.**
Box plot showing a titration of starvation in the Grooming vs Feeding assay.
Colored bars correspond to different p values
n= 10

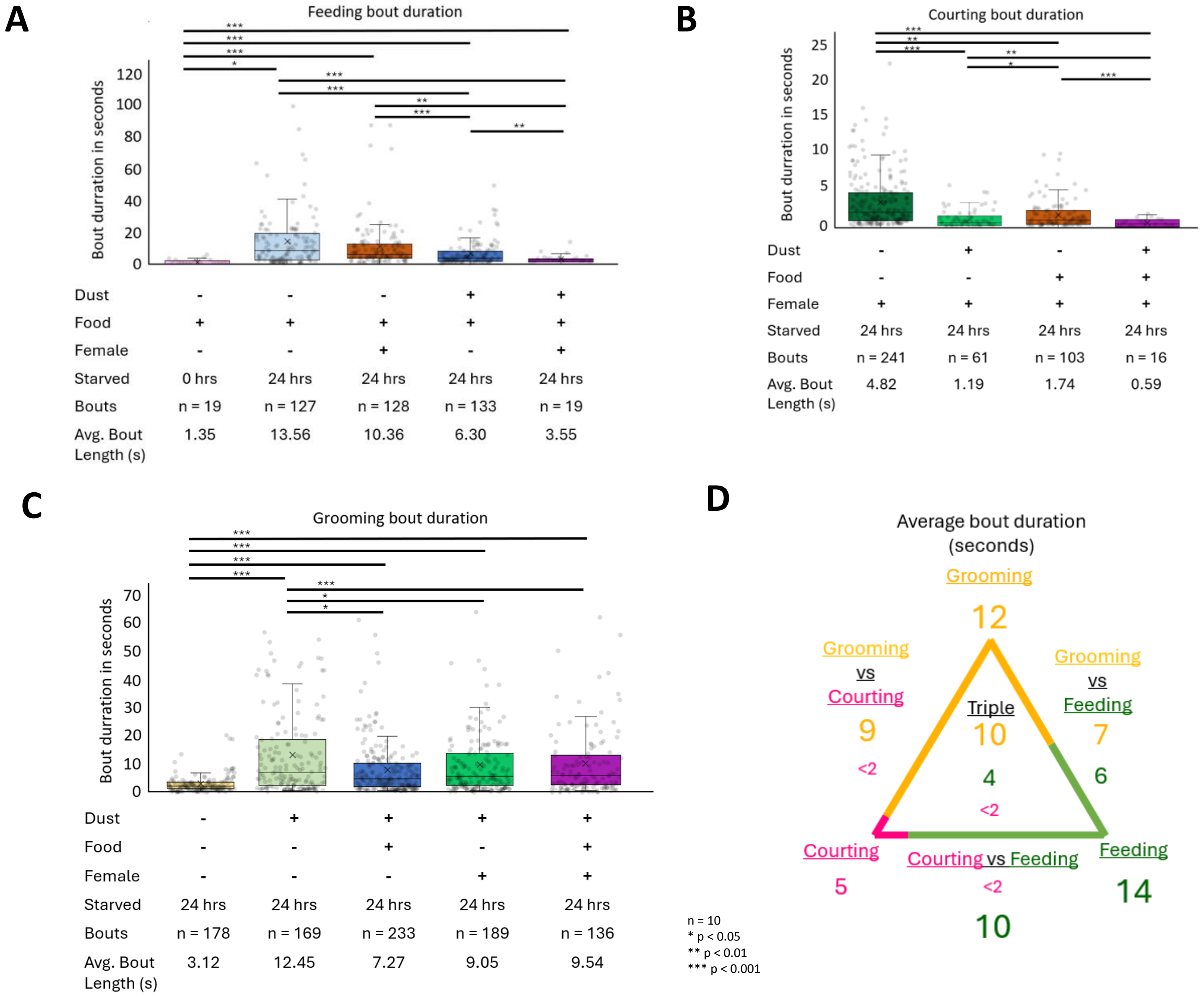
A
Feeding bout duration
Bout durration in seconds
0 20 40 60 80 100 120
*** *** *** * *** *** ** *** **
Dust - - - + +
Food + + + + +
Female - - + - +
Starved 0 hrs 24 hrs 24 hrs 24 hrs 24 hrs
Bouts n = 19 n = 127 n = 128 n = 133 n = 19
Avg. Bout Length (s) 1.35 13.56 10.36 6.30 3.55
B
Courting bout duration
Bout durration in seconds
0 5 10 15 20 25
*** ** *** ** * ***
Dust - + - +
Food - - + +
Female + + + +
Starved 24 hrs 24 hrs 24 hrs 24 hrs
Bouts n = 241 n = 61 n = 103 n = 16
Avg. Bout Length (s) 4.82 1.19 1.74 0.59
C
Grooming bout duration
Bout durration in seconds
0 10 20 30 40 50 60 70
*** *** *** *** *** * *
Dust - + + + +
Food - - + - +
Female - - - + +
Starved 24 hrs 24 hrs 24 hrs 24 hrs 24 hrs
Bouts n = 178 n = 169 n = 233 n = 189 n = 136
Avg. Bout Length (s) 3.12 12.45 7.27 9.05 9.54
D
Average bout duration (seconds)
Grooming
12
Grooming vs Courting
9
<2
Grooming vs Feeding
7
6
Triple
10
4
<2
Courting
5
Courting vs Feeding
<2
10
Feeding
14
n = 10
* p < 0.05
** p < 0.01
*** p < 0.001

**Supplemental 2: Effect of conflicting choice on bouts of behavior.**
A) A box plot with a dot plot overlay depicting individual feeding bout lengths from ten flies each within five minutes for each condition. B) A box plot with dot plot overlay depicting courting bouts (excluding orienting). C) a box plot with dot plot overlay depicting grooming bouts. Outliers are not depicted. D) A summary triangle showing average bout duration in seconds under the dusted and starved (Grooming); food and starved (Feeding); virgin female and starved (Courting); dusted, starved, and food (Grooming vs Feeding); dusted, starved, and virgin female (Grooming vs Courting); starved, food, and virgin female (Courting vs Feeding); and starved, dusted, food, and virgin female (Triple) conditions.

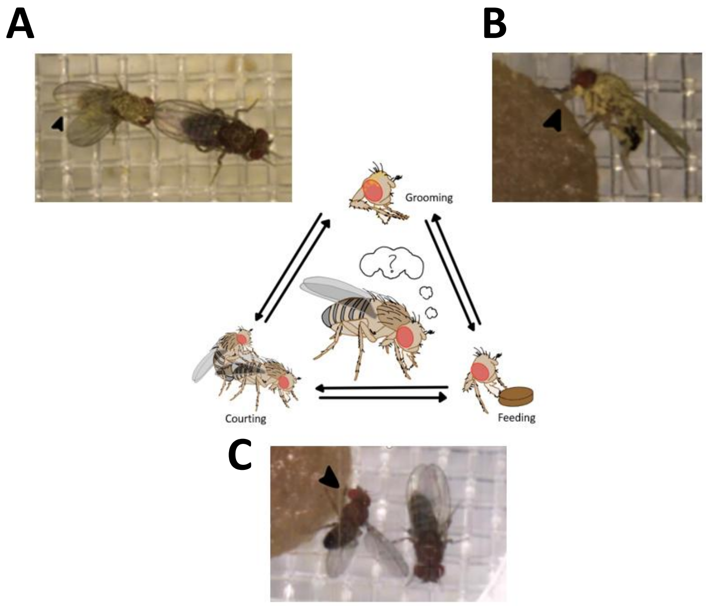
A
B
Grooming
Courting
Feeding
C

**Supplemental 3: Simultaneous behaviors in competition assays.**
A) A male fly simultaneously sing singing and back leg grooming (arrow). B) A male fly simultaneously feeding (arrow) and back leg grooming. C) A male fly simultaneously feeding (arrow) and wing singing

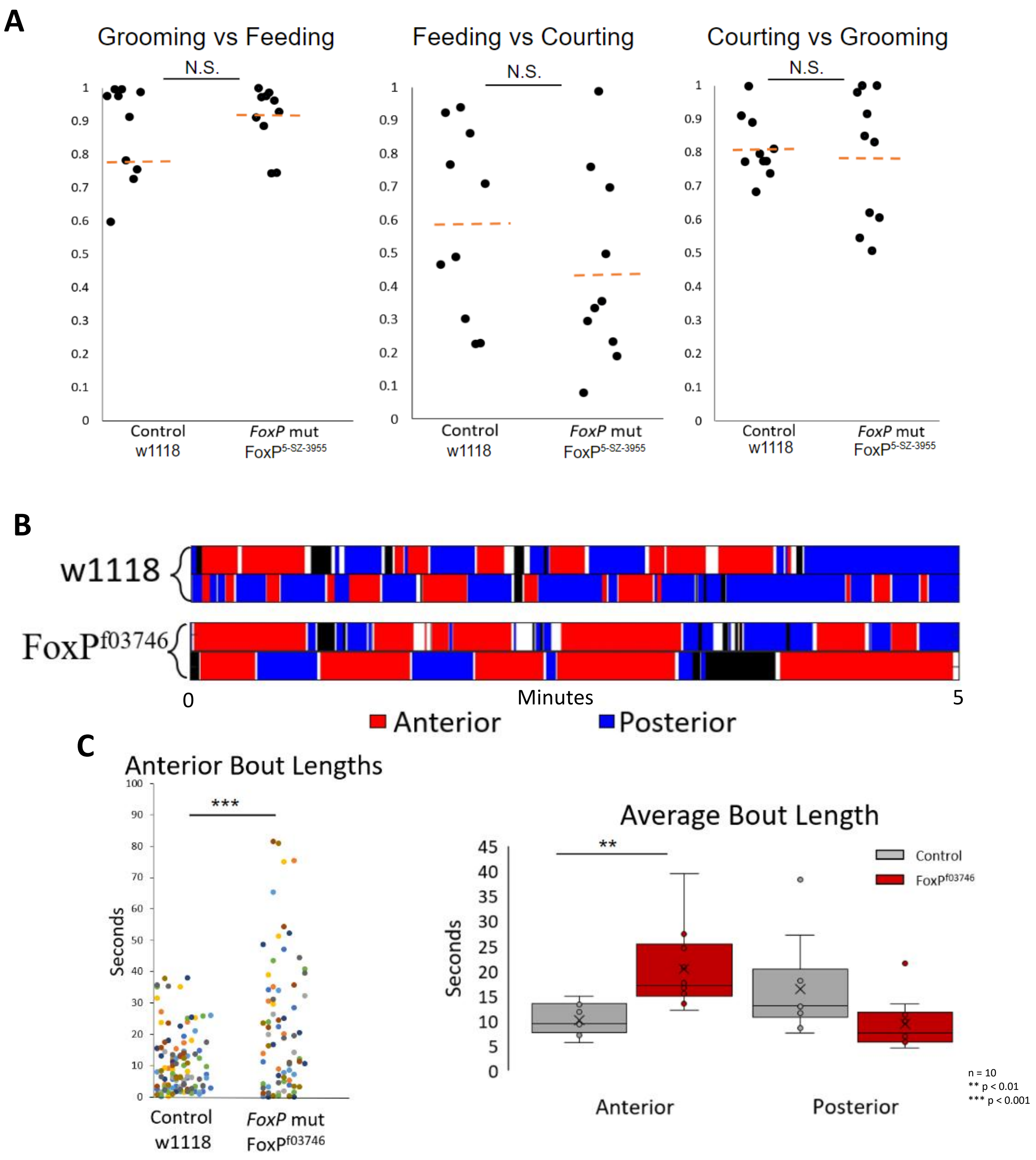
A
Grooming vs Feeding
Feeding vs Courting
Courting vs Grooming
N.S.
Control w1118
FoxP mut FoxP5-SZ-3955
B
w1118
FoxPf03746
0
Minutes
5
Anterior
Posterior
C
Anterior Bout Lengths
***
Seconds
Control w1118
FoxP mut FoxPf03746
Average Bout Length
**
Control
FoxPf03746
Anterior
Posterior
n = 10
** p < 0.01
*** p < 0.001

**Supplemental 4: *FoxP* mutants show a change in type of grooming behaviors compared to a w1118 control.**

A) A preference index of a *FoxP* mutant during the three competition assays. B) A sample ethogram of *FoxP* mutants grooming over a five minute time window looking at anterior grooming (red), posterior grooming (blue), walking (black), and standing (white). C) A dot plot looking at anterior grooming bouts in a five minute period with dusted flies. D) A box plot comparing anterior and posterior grooming in dusted flies

n = 10

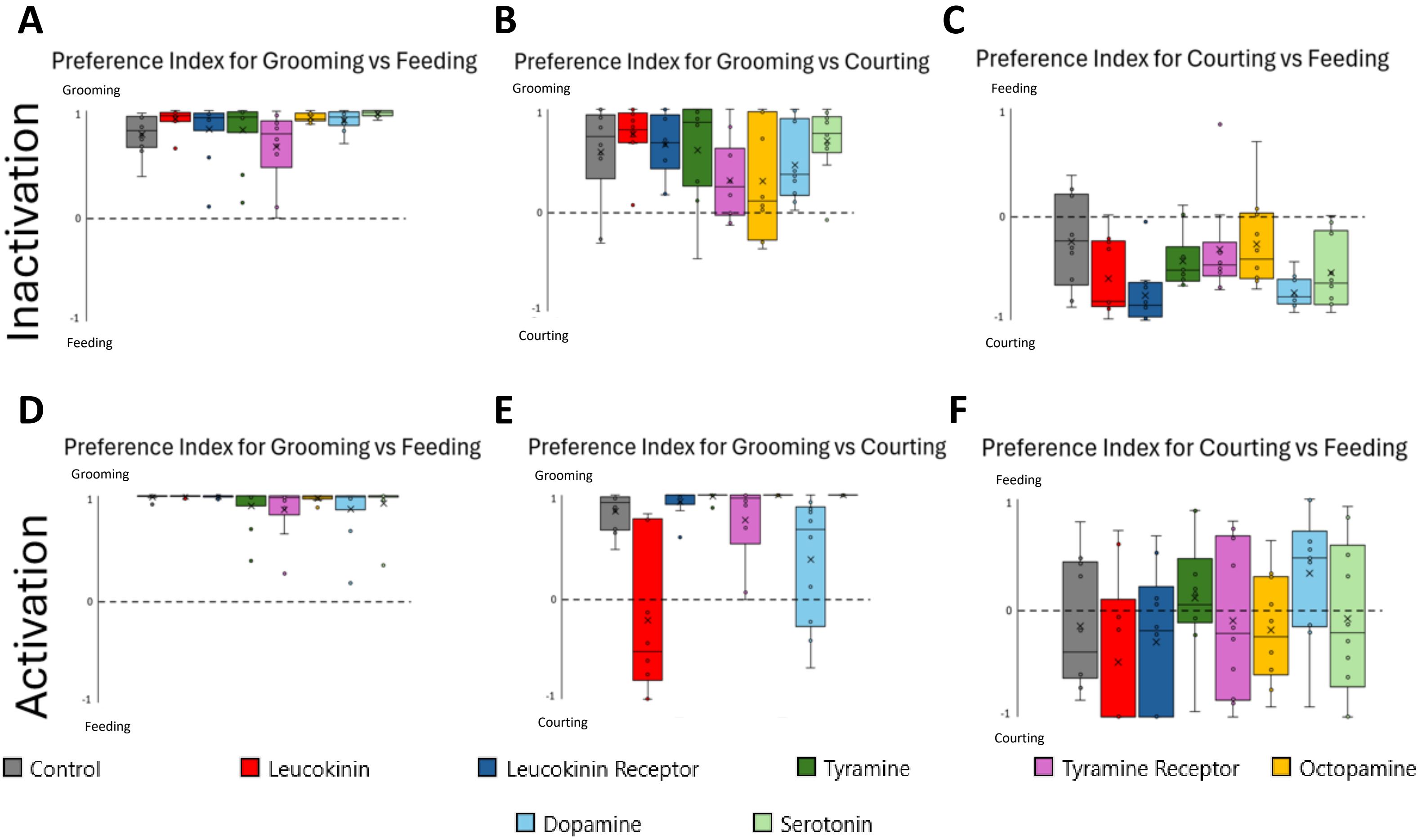
A
Preference Index for Grooming vs Feeding
Grooming
Feeding
B
Preference Index for Grooming vs Courting
Grooming
Courting
C
Preference Index for Courting vs Feeding
Feeding
Courting
Inactivation
D
Preference Index for Grooming vs Feeding
Grooming
Feeding
E
Preference Index for Grooming vs Courting
Grooming
Courting
F
Preference Index for Courting vs Feeding
Feeding
Courting
Activation
Control
Leucokinin
Leucokinin Receptor
Tyramine
Tyramine Receptor
Octopamine
Dopamine
Serotonin

**Supplemental 5: Screen results of activating and inactivating neurons tagged for different proteins.**

A) Preference index for Grooming vs Feeding when inactivating screened neurons subsets. B) Preference index for Grooming vs Courting when inactivating screened neurons subsets. C) Preference index for Courting vs Feeding when inactivating screened neurons subsets. D) Preference index for Grooming vs Feeding when activating screened neurons subsets. E) Preference index for Grooming vs Courting when activating screened neurons subsets. F) Preference index for Courting vs Feeding when activating screened neurons subsets.

n = 10

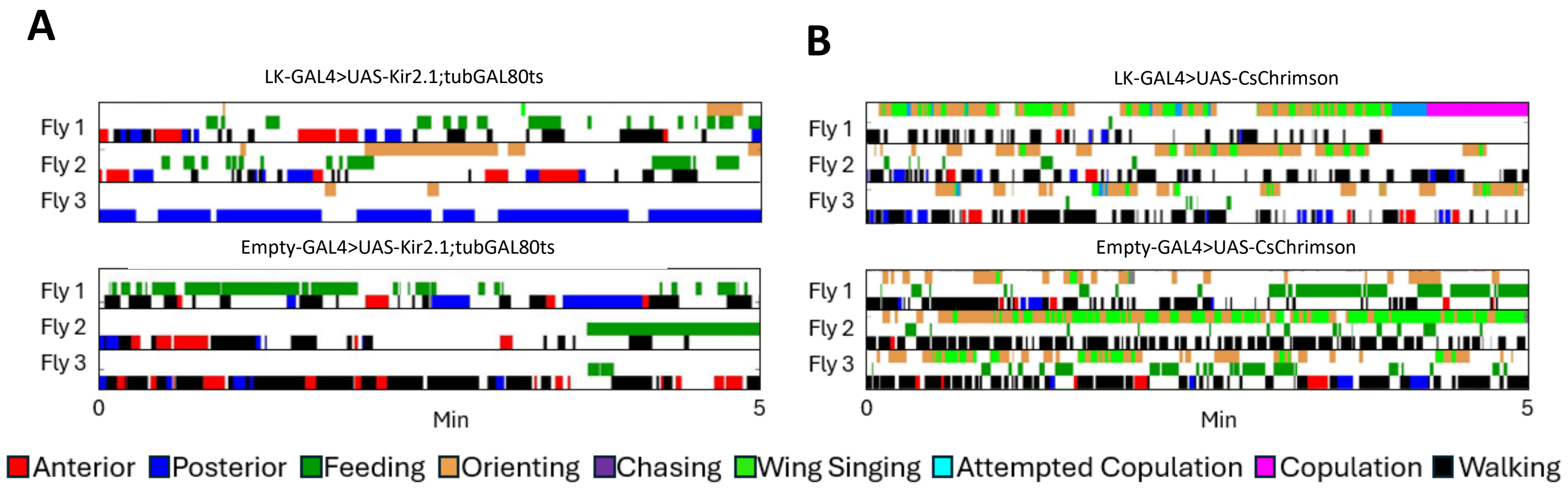
A
LK-GAL4>UAS-Kir2.1;tubGAL80ts
Fly 1
Fly 2
Fly 3
Empty-GAL4>UAS-Kir2.1;tubGAL80ts
Fly 1
Fly 2
Fly 3
0
Min
5
B
LK-GAL4>UAS-CsChrimson
Fly 1
Fly 2
Fly 3
Empty-GAL4>UAS-CsChrimson
Fly 1
Fly 2
Fly 3
0
Min
5
Anterior
Posterior
Feeding
Orienting
Chasing
Wing Singing
Attempted Copulation
Copulation
Walking

**Supplemental 6: Examples of ethogram analysis of individual components of the grooming, feeding, and courting behaviors.**

A) Ethograms depicting Feeding vs Courting behavior when inactivating leukokinin producing neurons. Grooming and courting behaviors are broken into micro-behaviors. B ) Ethograms depicting Feeding vs Courting behavior when activating leukokinin producing neurons

n = 10
